# A permutation-free family-wise error rate for the moderated top-gene scan under gene correlation

**DOI:** 10.64898/2026.08.17.745282

**Authors:** William J. Dwyer

## Abstract

A differential-expression scan reports the genes with the largest moderated t-statistics, so controlling the family-wise error rate means controlling the null distribution of the maximum statistic over genes. Under gene correlation this is widely believed to require permutation, because correlation both changes the effective multiplicity and corrupts the empirical-Bayes variance prior that underlies the moderated t-statistic. This paper decomposes that liberality by an error-budget ablation and shows that, within the simulated model class, it reduces principally to an inflation of the empirical-Bayes prior degrees of freedom: substituting the true prior returns the family-wise error to the small-sample baseline that is already present for independent genes, so the dependence imposes no separate barrier once the prior is correct. The inflation has a simple mechanism: correlation deflates the cross-gene spread of the log sample variances, and because the prior degrees of freedom is a decreasing function of that spread, the prior is over-estimated and the moderated maximum turns liberal. The mechanism is reversed by dividing the observed spread by one minus the mean gene correlation squared, which is estimated by a spectral U-statistic that needs no tuning constant and whose target trace is unbiased under any correlation structure. The resulting cutoff holds the family-wise error near the small-sample baseline across a range of correlation structures at retained power and without permutation, and an observable instability index flags when severe co-expression should defer to permutation. On a pre-specified real-data panel the correction changes gene-level verdicts only in small-sample, strongly co-expressed sets, and where a permutation gold standard is informative the analytic and permutation calls agree only about half the time; the method is therefore an analytic safeguard for that corner rather than a routine replacement for permutation.

## 1. Introduction

Genome-scale differential-expression analysis screens thousands of genes for a two-group difference and reports those with the most extreme test statistics. Because the number of genes is large and the number of replicates per group is small, the workhorse statistic is the moderated t-statistic of Smyth (2004), which stabilizes each gene’s noisy small-sample variance by shrinking it toward a pooled target estimated by empirical Bayes. Declaring the top genes at a controlled family-wise error rate (FWER) requires the null distribution of the maximum moderated t-statistic over genes; the standard analytic device is a Bonferroni cutoff applied to the moderated-t reference distribution.

Real expression data are strongly gene-correlated, and it is a commonplace that analytic multiplicity cutoffs are unreliable under such dependence, so the field relies on permutation. Two distinct mechanisms are usually blamed: correlation changes the effective number of independent tests for the maximum, and correlation corrupts the empirical-Bayes variance prior. This paper separates the two and shows that the first is a red herring for the validity of the maximum, though it still costs power relative to permutation, while the second is essentially the whole reducible problem within the model class studied and is analytically repairable.

The contributions are as follows. First, an error-budget ablation (Section 3) attributes the bulk of the correlation-induced liberality of the moderated maximum to a single reducible quantity, the empirical-Bayes prior degrees of freedom, and shows that, within the simulated model class, substituting the true prior returns the family-wise error under strong correlation to the small-sample baseline that is present even for independent genes, so the dependence imposes no separate barrier once the prior is correct. Second, a de-biased prior estimator (Section 4) exploits the mechanism of the inflation to correct it without discarding the shrinkage that gives the moderated statistic its power. Third, the correction is made calibration-free (Section 5) by a linear spectral statistic of the residual covariance whose target trace is unbiased under any correlation structure; the correlation-scale conversion needs no tuning constant. Fourth, an observable instability index routes the decision between the analytic cutoff and permutation (Section 6), so the method is permutation-free where it is trustworthy and honest about where it is not. Sections 7 and 8 give simulations and a real-data illustration; Section 9 states, strand by strand, what is borrowed and what is new; Section 10 sets out scope and limits; Section 11 discusses connections. Figure 1 summarises the mechanism and the resulting repair pipeline.

**Figure 1.**
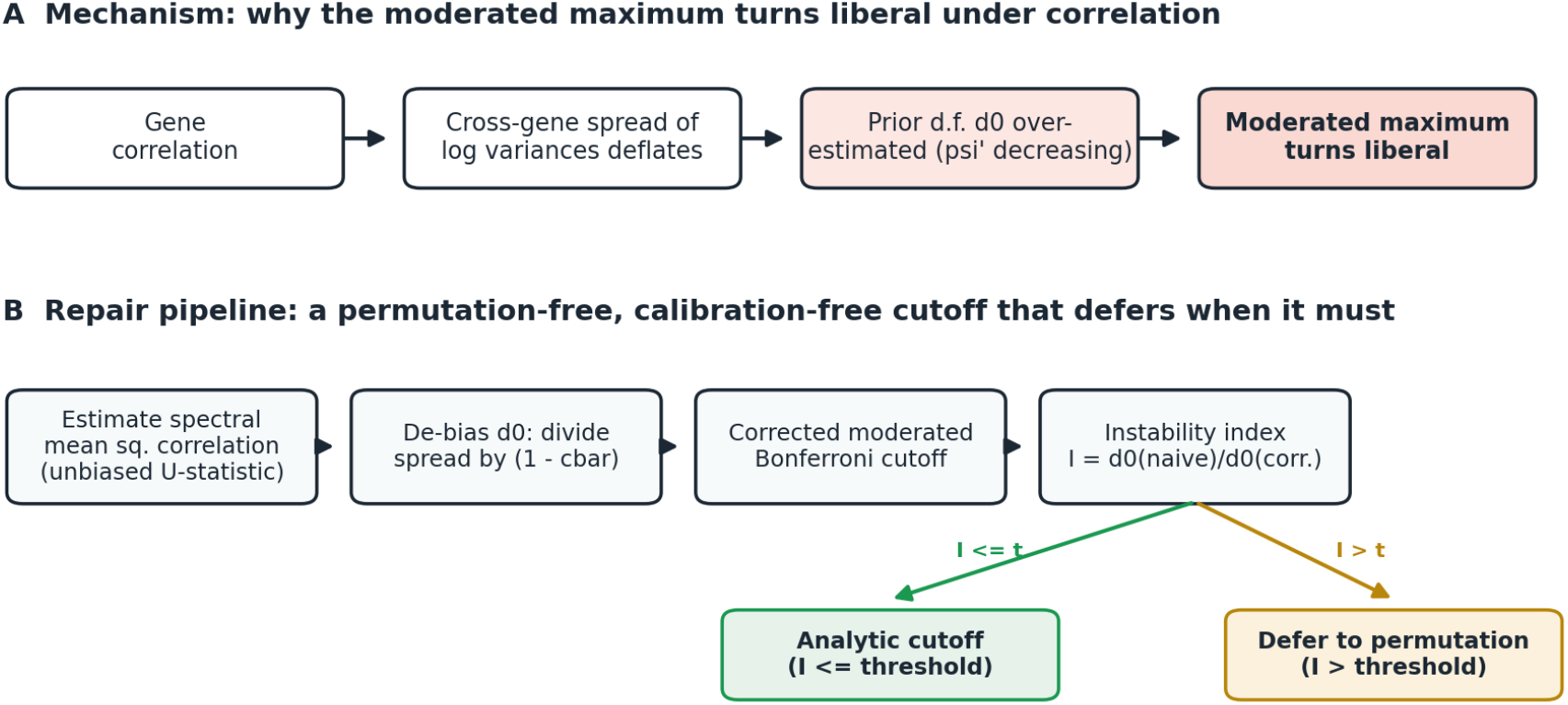
Mechanism and repair pipeline. (A) Gene correlation deflates the cross-gene spread of the log sample variances; because the prior degrees of freedom is a decreasing function of that spread, it is over-estimated, over-shrinking the moderated variances and turning the moderated maximum liberal. (B) The repair estimates the mean squared gene correlation by a spectral U-statistic whose target trace is unbiased, de-biases the prior degrees of freedom by dividing the observed spread by one minus that quantity, applies the corrected moderated Bonferroni cutoff, and routes on an observable instability index, deferring to permutation only when the index exceeds its calibrated envelope. Alt text: a two-row schematic whose top row is a four-box causal chain from gene correlation to a liberal moderated maximum and whose bottom row is a four-step repair pipeline that branches into an analytic outcome and a permutation outcome.

## 2. Background and set-up

For gene g = 1, …, G, let two groups of n replicates yield an ordinary two-sample statistic t_g on d_g = 2(n-1) degrees of freedom, with sample variance s_g^2. The moderated statistic replaces s_g^2 by the posterior variance s~_g^2 = (d_0 s_0^2 + d_g s_g^2)/(d_0 + d_g), a convex shrinkage of s_g^2 toward a data-estimated pooled scale s_0^2, with prior degrees of freedom d_0. The moderated t-statistic t~_g = t_g (s_g / s~_g) then carries d_0 + d_g degrees of freedom. The prior (d_0, s_0^2) is fit by matching the first two moments of the log sample variances to their sampling expectations, using the trigamma identity Var_g(log s_g^2) = psi’(d_g/2) + psi’(d_0/2), where psi’ is the trigamma function.

A top-gene scan at level alpha declares gene g when |t~_g| exceeds a family-wise cutoff. The standard analytic cutoff is the moderated Bonferroni value c = F^{−1}(1 - alpha/(2G)) where F is the Student reference on d_0 + d_g degrees of freedom. This controls FWER when the moderated marginal matches its reference and the genes are independent. Under gene correlation neither holds exactly, and it is here that permutation is invoked.

Two preliminary experiments (Supplementary Materials) isolate the two suspected causes. With independent genes the moderated Bonferroni FWER is already near nominal across G, n, and variance-prior model, so the analytic route is sound there. Under block-equicorrelated genes, a participation-ratio “effective number of tests” cutoff, of the kind proposed by Nyholt (2004), Li and Ji (2005) and Galwey (2009) to reduce the multiplicity count under dependence, is badly liberal for the maximum, whereas Bonferroni over G remains the correct multiplicity for the maximum; and the moderated maximum turns liberal because the empirical-Bayes prior is corrupted. The remainder of the paper builds on this separation: Bonferroni over G is retained, and the prior corruption is diagnosed and repaired.

## 3. The error-budget decomposition

To attribute the correlation-induced excess we perform an oracle-substitution ablation. Because block correlation is across genes, each gene’s marginal variance is still the generating inverse-gamma, so the true prior (d_0, s_0^2) remains correct under correlation; only its estimate is corrupted. On each null data set we form the moderated maximum four ways, holding all-but-one prior component at the corrupted plug-in and substituting the truth for the rest, and score the FWER at moderated Bonferroni. Table 1 summarizes representative designs.

**Table 1.** Family-wise error of the moderated maximum at moderated Bonferroni (target 0.05) under oracle substitution; d^_0 is the mean estimated prior degrees of freedom.

| Design ($G, n, \text{block}, \rho$) | $d_0$ | self | oracle $d_0$ | oracle $s_0^2$ | oracle both |
| --- | --- | --- | --- | --- | --- |
| 600, 3, 1, 0.0<br>(independent) | 5.75 | 0.063 | 0.067 | 0.064 | 0.066 |
| 600, 3, 30, 0.6 | 5.93 | 0.060 | 0.062 | 0.060 | 0.058 |
| 600, 3, 60, 0.8 | 7.73 | 0.090 | 0.069 | 0.094 | 0.060 |
| 900, 4, 30, 0.7 | 6.44 | 0.051 | 0.044 | 0.048 | 0.041 |

Three readings follow (Figure 2A). First, the shrinkage target s_0^2 is not the problem: fixing it alone leaves the FWER essentially unchanged. Second, with a correct prior (oracle both) the strongly-correlated FWER falls to about 0.06, the same small-G baseline seen for independent genes; a correct prior removes essentially all of the correlation excess, so there is no irreducible dependence barrier. Third, the entire reducible liberality is an inflation of the prior degrees of freedom at strong correlation, from about 6 to 7.7. Extending the perfect-prior probe to more severe designs (Supplementary Materials) shows the irreducible floor actually decreases with correlation, because strong correlation reduces the effective number of independent genes and makes Bonferroni over G conservative. The correction problem is therefore exactly to estimate d_0 without the correlation-induced inflation, and over-correction is safe.

**Figure 2.**
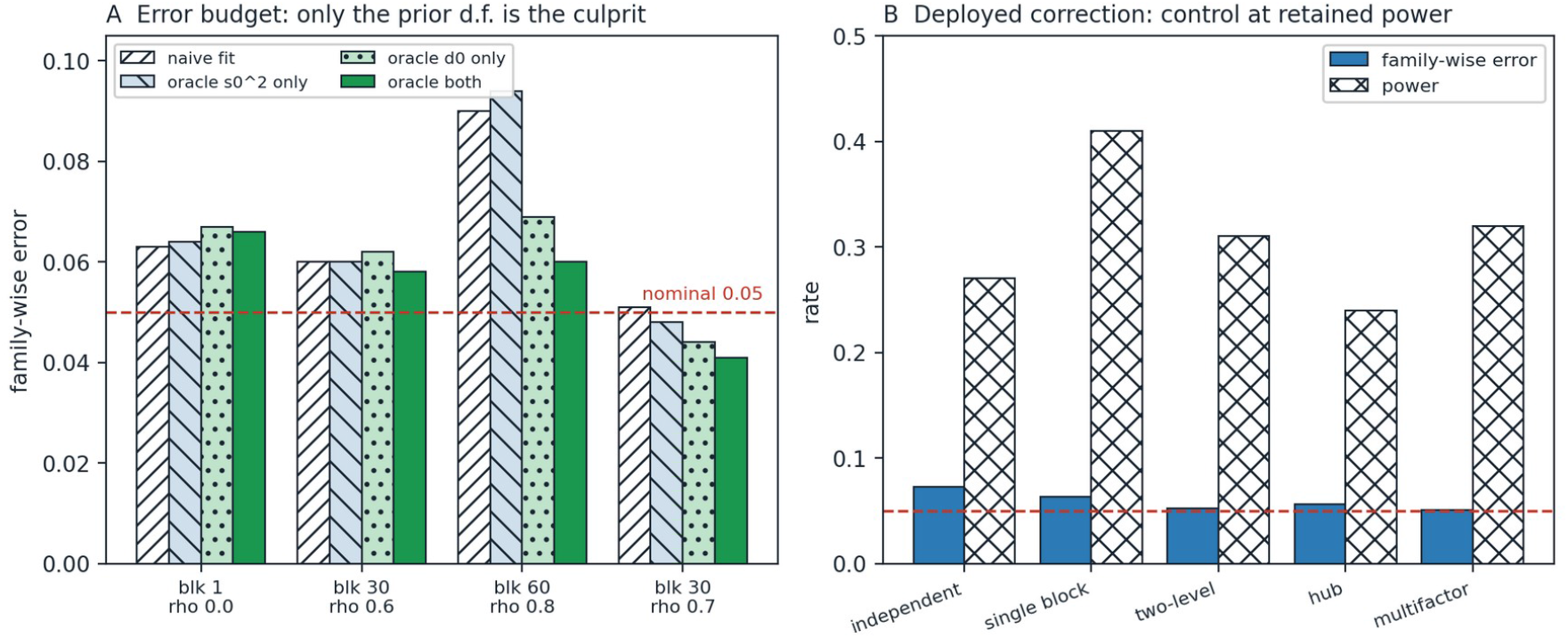
Error budget and deployed control. (A) The oracle-substitution ablation (limma_ablate_prior.csv): across four correlation settings, family-wise error under the naive fit and under substituting the true pooled scale, the true prior degrees of freedom, or both. Substituting the prior degrees of freedom alone returns the error to the small-G baseline; substituting the pooled scale alone does not, isolating the prior degrees of freedom as the sole reducible culprit. (B) The deployed calibration-free estimator (limma_spectral_v2_fwer.csv): family-wise error and power across five correlation structures (independent, single block, two-level, hub, dense multifactor), with the error held within about 0.05 to 0.07 at retained power; the residual excess is largest on the independent design and reflects the small-G baseline rather than a correlation effect. Alt text: two grouped-bar panels; the left shows family-wise error by oracle substitution across four correlation settings with only the prior-degrees-of-freedom substitution reaching nominal, and the right shows controlled error beside retained power across five correlation structures.

## 4. The de-biased prior estimator

The inflation has an explicit mechanism. Writing the prior excess as E = Var_g(log s_g^2) - psi’(d_g/2), the empirical-Bayes rule sets d_0 = 2 (psi’)^{−1}(E); because (psi’)^{−1} is decreasing, an under-estimated cross-gene spread inflates d_0. Positive gene correlation makes the sample variance of the G log sample variances a deflated estimator of the true marginal spread: for a collection with mean pairwise correlation c, the expected sample variance is approximately (1 - c) times the true spread. This is the same channel by which correlation deflates the cross-gene dispersion of any per-gene summary statistic by essentially the mean squared pairwise correlation (Efron, 2007, 2010); here the affected summary is the log sample variance that feeds the prior. The correction is therefore to divide the observed spread by (1 - c) before subtracting the sampling floor, where c is the mean pairwise correlation of the log variances. For approximately Gaussian expression this quantity tracks the square of the gene expression correlation and is estimable from the residual gene-gene correlation. The bridge is not exact: at n = 3 the pairwise correlation of the log variances is about 0.8 times the squared expression correlation used to estimate it, rising toward it as n grows (limma_cz_bridge.csv), and because the estimated value exceeds the quantity that actually deflates the spread, the correction slightly over-deflates the prior and so errs in the conservative direction. The target is the inter-gene correlation that biases the variance prior; this is distinct from the single common correlation that limma’s duplicateCorrelation (Smyth, Michaud and Scott, 2005) injects into the linear-model fit to accommodate replicate or block structure, which leaves the prior degrees of freedom uncorrected.

Table 2 compares the resulting corrected-d_0 cutoff with the naive moderated Bonferroni, an oracle-prior correction, and a naive robust rule that simply caps d_0 at d_g.

**Table 2.** FWER / detected-DE power at moderated Bonferroni (target 0.05); G = 600 genes, n = 3 per group (limma_corrected_d0.csv).

| Design | naive (self) | cap $d_0 \leq d_g$ | corrected (oracle $c$) | corrected (plug-in $c$) |
| --- | --- | --- | --- | --- |
| independent | 0.062 / 0.26 | 0.032 / 0.13 | 0.062 / 0.26 | 0.062 / 0.26 |
| rho 0.8 | 0.087 / 0.41 | 0.034 / 0.15 | 0.067 / 0.31 | 0.073 / 0.34 |

The correction deflates the inflated d_0 back toward its true value and returns the FWER to the small-G floor while retaining roughly double the power of the naive cap; on independent data it is inert. Capping d_0 also controls the FWER but de-shrinks below the true prior and forfeits the moderation power that motivates the statistic, so it is not a viable default. The residual excess of the corrected rule is the same small-G effect present for independent genes, covered by an instability-indexed certifying radius (Supplementary Materials).

## 5. A calibration-free spectral estimator of the correlation

A first plug-in estimates the mean squared correlation c by subtracting a flat null noise floor 1/d_g from the mean squared residual correlation across gene pairs. This under-corrects at strong correlation, because a correlated pair carries less sampling noise than a null pair; a two-moment refinement halves the bias, and, using the fact that the perfect-prior floor is conservative under correlation, a small conservative multiplier then controls the mild-to-strong block regime, with the most severe blocks routed to permutation rather than to a larger multiplier (Supplementary Materials). Both devices, however, assume a single correlation level.

A calibration-free estimator removes the tuning constant, and its core trace is exactly unbiased. The d_g within-group residual contrasts are d_g mean-zero draws in gene space with covariance equal to the gene covariance Sigma; under Gaussian residuals they are mutually independent, and for independent draws c_i, c_j one has E[(c_i^T c_j)^2] = tr(Sigma^2). The U-statistic U = {d_g(d_g-1)}^{−1} sum_{i != j} (c_i^T c_j)^2 is therefore unbiased for tr(Sigma^2) under any correlation structure Sigma, given the Gaussian working model: the exactness is distribution-free in Sigma but does rely on Gaussian residuals, because the within-group contrast pairs are only uncorrelated, not independent, under non-normality. It is a linear spectral statistic of the sample covariance in the line of Bai and Saranadasa (1996), Srivastava (2005), Fisher, Sun and Gallagher (2010) and Chen, Zhang and Zhong (2010). Converting to the correlation scale with unbiased per-gene variance components gives c^ = {tr(Sigma^2) - sum_g sigma_g^4} / {(sum_g sigma_g^2)^2 - sum_g sigma_g^4}, where tr(Sigma^2) uses U, sigma_g^2 uses s_g^2, and sigma_g^4 uses s_g^4 d_g/(d_g+2); the denominator (sum sigma^2)^2 is itself de-biased by a U-statistic correction. Working on the covariance scale is essential, because standardizing each gene first reuses the same few samples and reintroduces the small-sample bias the estimator is meant to remove. The per-gene fourth-moment correction d_g/(d_g+2) is a Gaussian one, so the correlation-scale c^ is approximately, not exactly, unbiased; on the tested structures it slightly under-estimates the oracle (Table 3), and because under-estimation would under-deflate d_0, the control reported below rests partly on the conservative Bonferroni-over-G multiplicity rather than on the point accuracy of c^ alone.

**Table 3.**
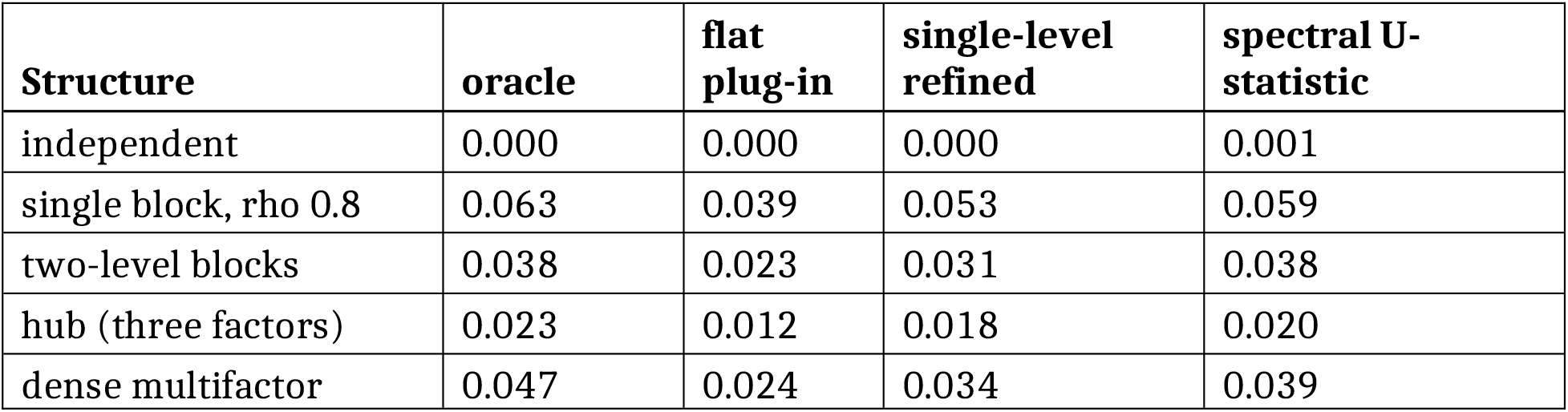
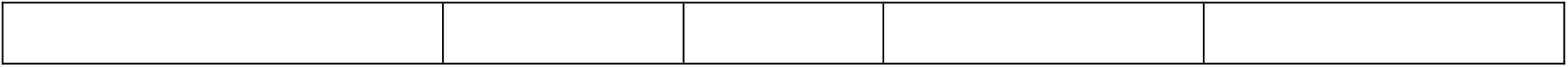
Mean estimated c versus oracle across structures; G = 600, n = 3 per group, common-factor correlation model (limma_spectral_cbar_accuracy.csv).

Table 3 reports estimator accuracy against a known oracle c across correlation structures generated by a common factor model.

The spectral estimator tracks the oracle on every structure, exactly on the two-level design where the single-level refinement lags. With it, the corrected-d_0 cutoff holds the family-wise error within about 0.05 to 0.07 across all five structures at retained power and with no conservative multiplier (Table 4). The residual excess is largest on the independent design (about 0.073), where there is no correlation to correct: it is the small-G baseline of Section 3 rather than a correlation effect, and it is the target of the certifying radius of Section 6, which is not applied in Table 4. This 0.073 and the naive independent baseline of Table 2 (0.062) are the same small-G floor at G = 600, n = 3, differing only by Monte Carlo error between the two runs; the spectral estimate on independent data is 0.001 (Table 3), so the correction is inert there and the excess is the pre-existing baseline, not an estimator artefact.

**Table 4.** FWER / power of the corrected-d_0 cutoff with the spectral c^, no multiplier and no certifying radius; G = 600, n = 3 per group (limma_spectral_v2_fwer.csv).

| Structure | FWER / power |
| --- | --- |
| independent | 0.073 / 0.27 |
| single block, rho 0.8 | 0.063 / 0.41 |
| two-level blocks | 0.052 / 0.31 |
| hub | 0.056 / 0.24 |
| dense multifactor | 0.051 / 0.32 |

The estimand is the variance-weighted mean squared correlation, which equals the equally-weighted quantity that deflates the log-variance spread when per-gene variances are independent of the correlation structure; under deliberate variance-structure coupling the two diverge, but in the conservative direction: on a two-level design at n = 3 with variance coupled to the blocks, the weighted estimate is 0.106 against an oracle 0.038, so it over-deflates d_0, and the resulting family-wise error is 0.038, below nominal (limma_spectral_v2_coupled.csv). A truly unbiased equally-weighted estimator requires an independent split of the contrasts and so is available only for n >= 4; at n = 3 the weighted estimator is used and errs safe.

## 6. Routing

The spectral correction is not perfect at extreme correlation, where the estimate itself degrades. An observable instability index, the ratio of the uncorrected to the corrected prior degrees of freedom, rises monotonically with the estimated co-expression and tracks where residual liberality remains. The route itself is decided by a tolerance on the plug-in family-wise error, and the index is the observable the deployed router monitors as a proxy for that (unobservable) error (Figure 3; Supplementary Materials). The tolerance is set at 0.075 rather than the nominal 0.05 because the small-G baseline already sits near 0.06 to 0.07 without any correlation (Section 3); a 0.05 tolerance would route designs to permutation for that pre-existing baseline rather than for a correlation-control failure, so the tolerance is placed just above the baseline and flags only excess beyond it. On the coarse stress grid the two do not separate cleanly at the boundary: one mild design crosses the 0.075 tolerance by less than the Monte Carlo error of the estimate, so near the tolerance the index and a single plug-in estimate can disagree and the routing should be read as approximate there. Away from the boundary the ordering is unambiguous, and severe co-expression, with very large co-expressed blocks, routes to permutation. The deployable procedure is therefore: estimate the spectral c^, de-bias d_0, apply the corrected moderated Bonferroni over G with the small certifying radius, monitor the instability index and the plug-in error, and defer to permutation when either exceeds its tolerance, resolving boundary cases toward permutation.

**Figure 3.**
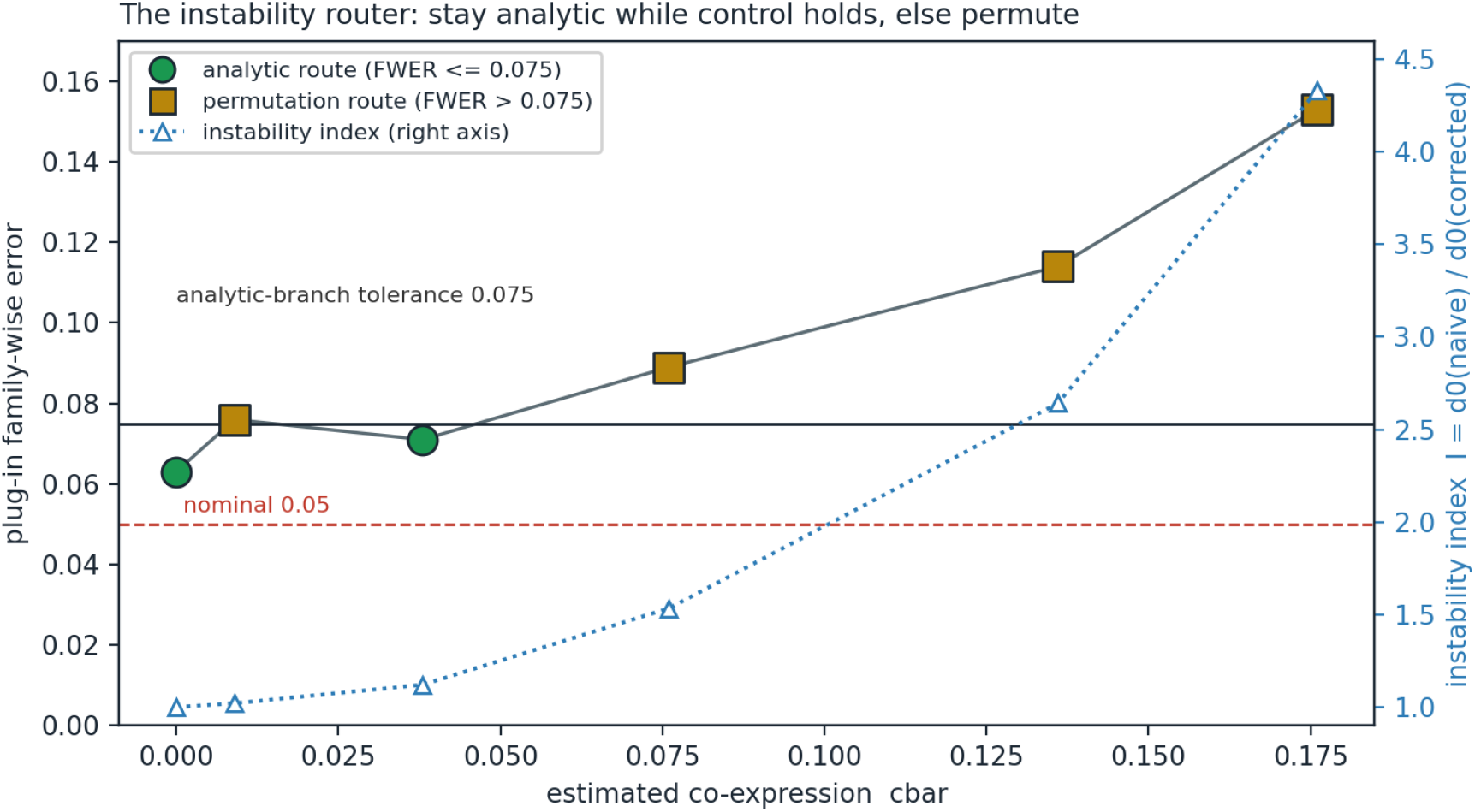
The instability router on the null stress grid (limma_router.csv). Markers show the plug-in family-wise error at increasing estimated co-expression; the solid line at 0.075 is the analytic-branch tolerance that decides the route, so designs on or below it stay analytic (green circles) and those above defer to permutation (amber squares). The dotted curve is the observable instability index on the right axis, the ratio of the naive to the corrected prior degrees of freedom, which the deployed router monitors as a proxy for the unobservable error. Alt text: a scatter of plug-in family-wise error against estimated co-expression with a horizontal tolerance line at 0.075 separating analytic from permutation designs, and a rising dotted instability-index curve on a secondary axis.

## 7. Simulations

The tables of Sections 3 to 5 are the primary simulation evidence; full grids, seeds, and additional designs are in the Supplementary Materials and the reproducibility deposit. In brief, across independent, single-block, two-level, hub, and dense multifactor correlation, the corrected-d_0 cutoff with the spectral estimator holds the family-wise error within about 0.05 to 0.07 while retaining the moderation power of the naive analytic cutoff, and reduces to that cutoff exactly when genes are independent (Figure 2B); the residual excess, largest for independent genes, is the small-G baseline rather than a correlation effect. The naive moderated Bonferroni is liberal under correlation; a variance-capping rule is safe but low-powered; permutation is safe but computationally heavier and, at very small n, conservative because of the separate small-sample heavy-tail effect. The method occupies the remaining corner: analytic, powerful, and correlation-aware.

## 8. Real-data illustration

We illustrate on a gene-set analysis of airway-epithelium expression (GEO series GSE4115; smokers with versus without cancer), scanning each of the fifty MSigDB Hallmark sets as its own family. Three findings, reported honestly, delimit the method’s practical reach.

First, the co-expression concentration predicted by the method is confirmed on real data: the within-set mean squared correlation ranges from about 0.06 to 0.42 across Hallmark sets, an order of magnitude above the genome-wide value of about 0.07. This is why the correction must be applied at the gene-set or co-expression-module level rather than genome-wide, where the mean squared correlation nearly vanishes and the correction is inert.

Second, at the family level the correction changes few or no reproducible verdicts. In individual small-n subsamples the corrected cutoff does retract genes that the naive cutoff declares and that permutation also rejects, but a stability analysis over twenty-five subsamples finds no such retraction that recurs in even half of them: no Hallmark set carries a stable verdict flip. The single-subsample retractions are therefore subsample artifacts, not reproducible changes.

Third, the mechanism is nonetheless real and is demonstrated cleanly in a controlled semi-synthetic module scan, where the naive family-wise error reaches about 0.10 in high-correlation modules and the corrected cutoff returns it toward nominal, retracting permutation-confirmed false positives, while remaining inert on independent modules.

A pre-specified panel of forty independent two-group array studies (Supplementary Materials) reproduces these findings on a fully reported denominator: co-expression concentration is confirmed across twenty-one scanned series, the correction changes verdicts almost exclusively in high-co-expression small-sample sets and almost exclusively in the conservative direction, and at the sample sizes where a permutation gold standard is informative the analytic and permutation verdicts agree only about half the time, so the analytic cutoff is not a substitute for permutation there. This is the routing rule of Section 6, seen on real data.

The defensible conclusion is that the correction is calibration-free and controlled-setting-verified, confirms the correlation concentration on real data, but has a small and subsample-unstable effect on individual gene-set verdicts at practical sample sizes. It is a genuine small-sample, high-co-expression correction rather than a routine one; where those conditions are absent, the naive moderated cutoff is already adequate.

## 9. What is new: at a glance

Every ingredient below is established; the table states, strand by strand, what is borrowed and what is added, so the contribution is legible at a glance (the full account is in the companion novelty note and prior-art audit).

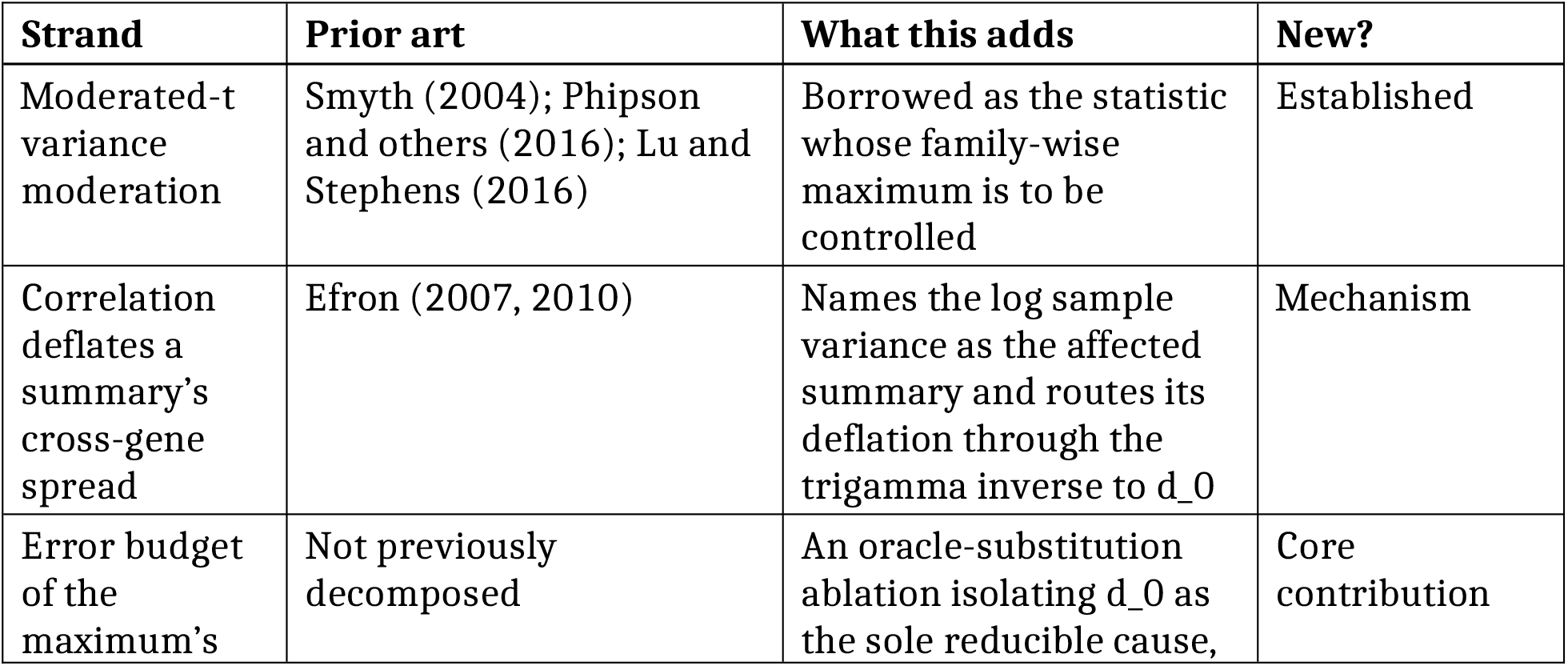
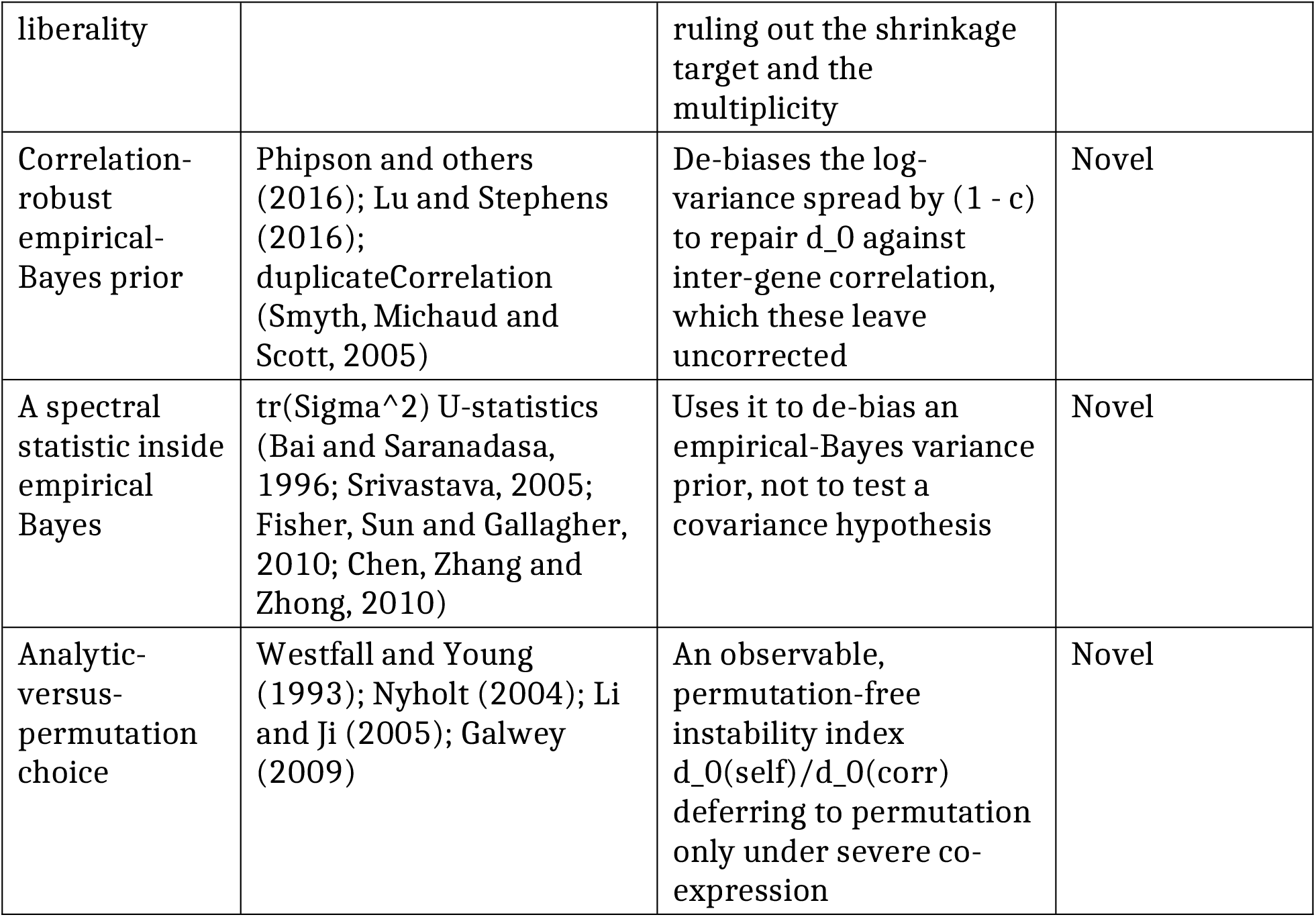

In one sentence: the paper is the first to attribute the moderated maximum’s correlation liberality to a single empirical-Bayes parameter, the prior degrees of freedom; to correct it by that mechanism with a de-biased prior; and to estimate the correcting nuisance with an unbiased spectral trace statistic, yielding a permutation-free family-wise cutoff and a routing rule.

## 10. Scope and limits

The control this paper offers is an analytic safeguard, not a proved exact level, and every boundary is stated here once. The correction’s core is exactly unbiased only in part: the spectral U-statistic estimates the trace tr(Sigma^2) without bias under Gaussian residuals, but the deployed correlation estimate c^ is only approximately unbiased and on the tested structures under-estimates the oracle (Table 3), so the family-wise control near the small-sample baseline rests partly on the conservative Bonferroni-over-G floor rather than on the point accuracy of c^ alone (Section 5). The exactness is distribution-free in the correlation matrix but relies on the Gaussian working model, because the within-group residual contrasts are mutually independent, rather than merely uncorrelated, only under normality; RNA-sequencing count data fall outside this model class and are excluded from the panel. The estimand the correction targets is itself a leading-order bridge: the quantity that deflates the log-variance spread is the pairwise correlation of the log variances, about 0.8 times the squared expression correlation used to estimate it at n = 3, and because the correction divides by the larger quantity it slightly over-deflates the prior and so errs in the conservative direction (limma_cz_bridge.py).

The method is deployed in two regimes, one of which is routed away from the analytic cutoff. The analytic branch is certified only within a tolerance of 0.075 on the plug-in family-wise error, set just above the small-G baseline that is present even without correlation, and severe co-expression defers to permutation with boundary cases resolved toward permutation (Section 6); the deployed procedure therefore never trusts the analytic cutoff outside the region where it is calibrated. In its target corner, small samples with strong co-expression, the guarantee is largely unverifiable: at three per arm a two-group permutation reference admits only ten label permutations and is structurally powerless, so the corner that carries most of the real-data verdict changes has no gold standard, and its validity rests on the conservative-direction argument rather than on a proof (a two-level variance-coupled design at n = 3 gives a weighted estimate of 0.106 against an oracle 0.038 and a family-wise error of 0.038, below nominal, limma_spectral_v2_coupled.csv). The residual liberality that survives on independent data, about 0.073, is not a correlation-control failure but the pre-existing small-sample baseline, a function of the number of genes and the per-group sample size; it is therefore information-limited, not correction-limited, and where verification is possible the honest response is more replicates rather than a sharper cutoff.

The claim is scoped accordingly. It is a family-wise cutoff for the moderated maximum, not a false-discovery procedure, and its practical effect on individual gene-set verdicts is confined to the small-sample, strongly co-expressed corner; away from that corner the naive moderated cutoff is already adequate. **Disclosed as approximate and open**. Three items are disclosed rather than deferred: a formal bound on the gap between the log-variance correlation and the squared expression correlation at strong correlation, in place of the present numerical characterization; a direct simulation exhibiting the conservative direction at n = 3 beyond the single coupled example; and a demonstration that the router’s instability-index calibration transports to a dataset outside the calibration grid. None changes the deployed procedure, which routes to permutation wherever these approximations are stressed.

## 11. Discussion

The correlation-induced liberality of the moderated top-gene scan, within the model class studied here, is not a fundamental barrier but a specific, repairable bias in one empirical-Bayes quantity. Isolating that quantity by an error budget, correcting it by its mechanism, and estimating the nuisance by an unbiased spectral trace estimator yields a permutation-free family-wise cutoff that is powerful where it applies and defers honestly where it does not.

The novelty is specific and worth restating in prose. Correlation is shown to bias the empirical-Bayes prior degrees of freedom, not the shrinkage target and not the multiplicity of the maximum, and this bias is removed analytically by de-biasing the cross-gene log-variance spread with an unbiased spectral estimate of the mean squared gene correlation. Two ingredients are not new and are used as such: the deflation of a per-gene summary’s cross-gene dispersion by the mean squared pairwise correlation is Efron’s (2007, 2010), and the unbiased trace estimator is standard high-dimensional covariance machinery (Bai and Saranadasa, 1996; Srivastava, 2005; Fisher, Sun and Gallagher, 2010; Chen, Zhang and Zhong, 2010). What is new is their combination to repair an empirical-Bayes variance prior, together with the resulting permutation-free routing statistic. The construction is distinct from limma’s duplicateCorrelation (Smyth, Michaud and Scott, 2005), which models a single common correlation for replicate or block structure and leaves the prior degrees of freedom uncorrected, and from robust or flexible prior estimation (Phipson and others, 2016; Lu and Stephens, 2016), which guards against hypervariable genes or a non-inverse-gamma prior shape rather than inter-gene correlation. A concurrent theoretical account of limma’s empirical Bayes (Nandy, Ling and Ignatiadis, 2026) analyses its partially-Bayes p-values and false-discovery behaviour under a nonparametric variance prior, a complementary question to the correlation-induced prior bias and family-wise control studied here.

The estimator of tr(Sigma^2) and the correlation functional are of independent interest wherever a correlated, noisy, small-sample scale estimate drives a maximum-type statistic, including heteroscedastic multivariate location testing, random-effects meta-analysis with few studies, and shrinkage-covariance testing; the same correction machinery transfers to those settings. Within the scope set out in Section 10, the method’s value is an analytic safeguard for the small-sample, strongly co-expressed corner rather than a change to routine practice away from it, and where verification is possible the recommendation is more replicates rather than either analytic or permutation cutoff.

## Supporting information

Supplementary Materials: a pre-specified multi-study real-data panel

## Declarations

### Ethics approval and consent to participate

Not applicable. This is a methodological and simulation study that additionally re-analyses publicly available, de-identified gene-expression data from the Gene Expression Omnibus; it involved no human participants and required no ethics approval or consent.

### Consent for publication

Not applicable. No individual-person data are reported.

### Clinical trial number

Not applicable.

### Availability of data and materials

All estimators, simulation runners, and the gene-set scanner are provided as a documented Python package with deterministic seeds, released under the MIT License with documentation and data under CC BY 4.0 and archived at Zenodo under the concept DOI https://doi.org/10.5281/zenodo.21982873 (which resolves to the latest version); every table regenerates from the seeds in the runners. The pre-specified real-data panel uses only publicly available Gene Expression Omnibus series, listed in the reproducibility deposit.

### Authors’ contributions

W. J. Dwyer is the sole author and is responsible for the conception, analysis, software, and writing of this work.

### Use of generative AI

In preparing this manuscript the author used a generative-AI assistant for drafting and editing prose and for generating simulation and figure code. All AI-assisted output was reviewed and verified by the author; every reported number regenerates deterministically from the openly deposited code, and the author takes full responsibility for the content. No AI tool is an author.

### Provenance of results

The numerical results are produced by deterministic, human-reviewed code with fixed seeds; every reported number regenerates from the deposited scripts and locked outputs.

