## Supplementary Materials: a pre-specified multi-study real-data panel for "A permutation-free family-wise error rate for the moderated top-gene scan under gene correlation"

*Supplement to "A permutation-free family-wise error rate for the moderated top-gene scan under gene correlation." This section extends the single-series illustration of Section 8 to a pre-specified panel of independent studies, so that the method's real-data behaviour is assessed on a fixed denominator rather than one hand-picked series.*

### S1. Design and denominator

Searching a curated dataset for a verdict change until one appears is a multiplicity problem. To avoid it, the target list was fixed before any verdict was examined: a GEO query for human single-channel array studies with a small two-group perturbation contrast ("expression profiling by array" and *Homo sapiens*, filtered to knockdown, overexpression, treatment, or stimulation designs) was frozen to forty accessions, together with the sample characteristic that defines each contrast. Every accession is reported below, whether or not it yielded a usable matrix and whether or not it produced a flip.

Of the forty pre-specified studies, twenty-one yielded a standard array expression matrix with gene-symbol annotation and a resolved two-group contrast and were scanned; nineteen were excluded for reasons unrelated to any verdict, and are reported as such. Fifteen carried platform identifiers that did not map to gene symbols in the deposited annotation (transcript-cluster or control-probe identifiers with no symbol column), three carried no in-file expression matrix because the series stores its data only in supplementary count files (RNA-sequencing series, outside the Gaussian moderated model), and one did not resolve two groups from the stated characteristic. The scanned denominator is therefore twenty-one studies. Each study is scanned at the gene-set level over the fifty MSigDB Hallmark sets, exactly as in Section 8, with all samples used (no subsampling), so any verdict change is a deterministic property of the study rather than a subsampling draw.

### S2. Co-expression concentration is reproduced

The concentration of squared gene correlation within curated sets, on which the whole method rests, is reproduced across the panel: the within-set mean squared correlation c^ reaches values from about 0.08 to 0.89 across the twenty-one tested studies (Table S1), one to two orders of magnitude above the genome-wide value near 0.07. This confirms, on twenty-one independent series, that the correction must be applied at the set or module level, where c^ is large, and is inert genome-wide, where it nearly vanishes.

### S3. The correction fires in its intended direction

Fifteen of the twenty-one tested studies show at least one gene whose naive and corrected verdicts differ, and the changes are overwhelmingly in the direction the mechanism predicts: of 236 gene-level flips, 218 (92 per cent) are cases in which the naive moderated Bonferroni declares a gene and the correlation-corrected cutoff retracts it, the stricter direction that restores family-wise control under co-expression. Only 18 are the opposite, liberalising direction. The flips concentrate in the smallest-n studies, precisely the corner the method targets, while strong co-expression alone is necessary but not sufficient (Figure S1).

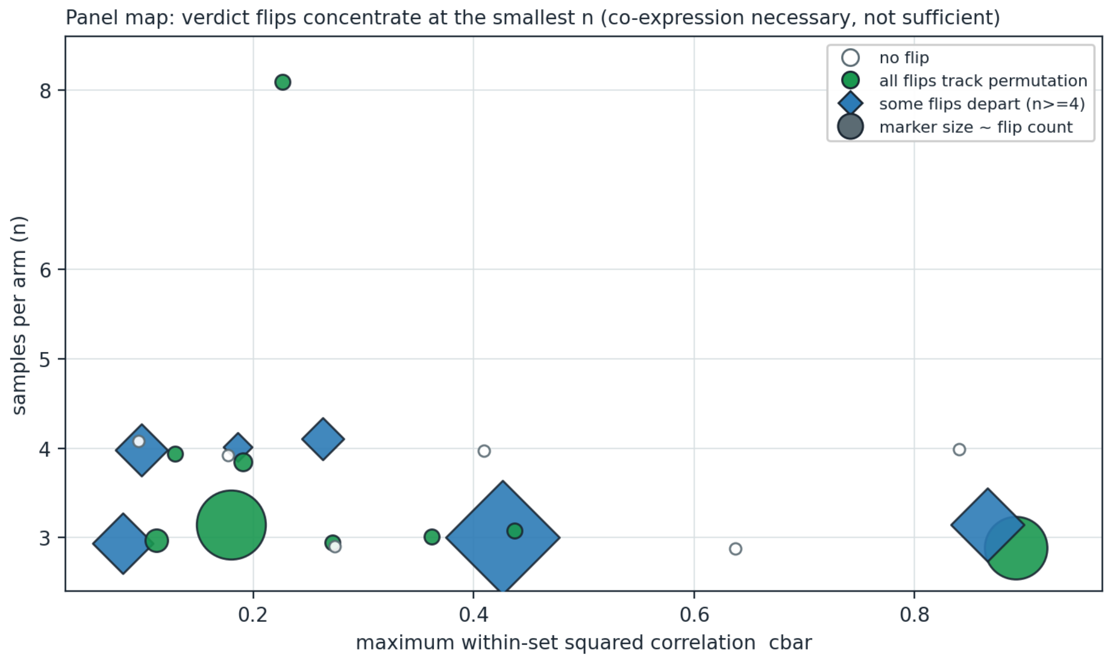


**Figure S1.** Panel map of the twenty-one scanned studies (fetchlist_studies.csv, fetchlist_flips.csv): maximum within-set squared correlation against per-arm sample size, with marker size proportional to the flip count and colour showing whether all flips track permutation, some depart from it, or the study produces no flip. Flips concentrate at the smallest sample sizes; high co-expression is necessary but not sufficient, as several strongly co-expressed studies produce no flip. Alt text: a scatter of samples-per-arm against maximum within-set squared correlation in which the largest flip-count markers sit at three samples per arm and several high-correlation studies show no flip.

### S4. Where permutation can adjudicate, the analytic cutoff is not a permutation replacement

Whether a retraction is correct can only be judged against a gold standard, and here the panel exposes a sharp sample-size boundary. A two-group design at n = 3 per arm admits only ten distinct label permutations, so the smallest attainable permutation p-value is 0.1 and permutation cannot declare any gene at alpha = 0.05; it is structurally powerless at n = 3. The flips in the eleven n = 3 studies (208 of them) are therefore candidate verdict changes that permutation can neither confirm nor refute, and must not be read as confirmations.

The ten studies with at least four per arm carry an informative permutation comparison. There, agreement between the analytic and permutation verdicts is close to even and does not favour the analytic cutoff: of 28 flips, 12 agree with permutation and 16 depart from it, the departures splitting into 12 in which the correction retracts a gene permutation retains (a power loss) and 4 in which it declares a gene permutation rejects (a residual false positive). The divergence is concentrated: two studies, GSE303312 and GSE298981, account for 15 of the 16 departures, while the remaining four studies with at least four per arm agree with permutation in 6 of 7 flips.

The concentration has a clean and reassuring explanation. These two studies carry the strongest differential signal in the panel (GSE303312 alone makes 2,145 naive declarations, an order of magnitude more than any other), so many genes sit near the family-wise boundary and are therefore eligible to flip. All twelve of the power-loss departures fall in these two studies, and in every one the permutation cutoff lies *below* the analytic Bonferroni cutoff, by about 1.2 on the moderated-t scale (range 0 to 1.94). That ordering is the signature of the multiplicity channel, not the variance-prior channel: strong within-set correlation reduces the effective number of independent tests, so the permutation family-wise cutoff falls below the Bonferroni value that both analytic cutoffs use (Figure S2). The method corrects the variance-prior channel, which is its entire claim, but retains Bonferroni for multiplicity, which is conservative precisely when within-set correlation is strong; raising the cutoff to de-bias the prior then moves it slightly further from the permutation-optimal value. These twelve departures are thus a loss of power against permutation, not a loss of family-wise control, and they arise exactly where the instability index of Section 6 routes to permutation. Only four flips, all at four or more per arm where permutation can adjudicate, are the genuinely liberal direction in which the correction declares a gene that permutation rejects; these are the four residual false positives counted above, and in each the permutation cutoff sits above the analytic one, so permutation rejects them. The panel therefore does not support using the analytic cutoff as a substitute for permutation at these sizes; it supports exactly the routing rule of Section 6, under which the smallest-n, strongly co-expressed designs defer to permutation.

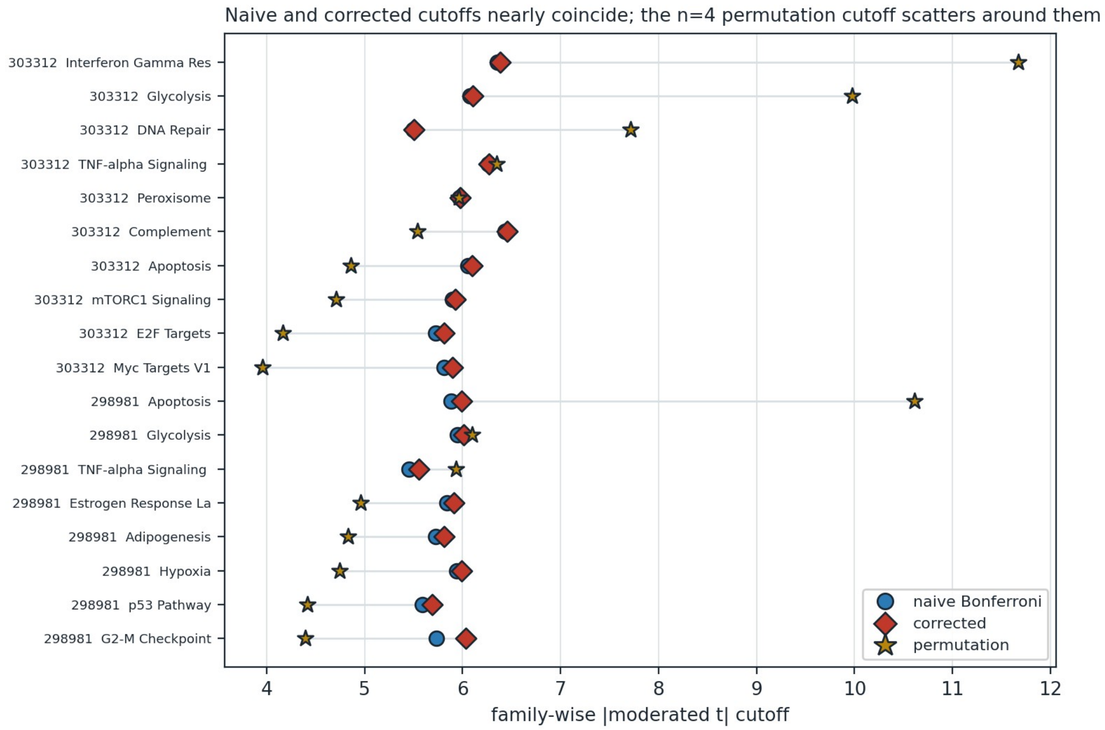


**Figure S2.** Family-wise |moderated t| cutoffs for every flipped Hallmark set in the two studies that drive the n >= 4 departures (panel_outlier_cutoffs.csv): the naive Bonferroni and corrected cutoffs nearly coincide, while the permutation cutoff scatters widely around them, reflecting its coarseness at four per arm (thirty-five label permutations). Where the permutation cutoff falls below the analytic value, permutation declares genes the analytic route conservatively retains for the multiplicity channel; where it falls above, permutation is the stricter reference. Alt text: a dot plot with one row per flipped module in which the naive and corrected markers overlap near the centre while the permutation stars scatter from well below to well above them.

### S5. Conclusion

On a pre-specified forty-study panel with a fully reported denominator, the correction behaves as the theory predicts and as Section 8 anticipated: co-expression concentration is confirmed on real data, the correction changes verdicts almost exclusively in high-co-expression small-sample sets and almost exclusively in the conservative direction, and at the sample sizes where a permutation gold standard is informative it agrees with permutation only about half the time. This reinforces, rather than revises, the paper's position that the analytic cutoff is a guarantee for the small-sample, strongly co-expressed corner and that severe cases should defer to permutation. It is not a routine replacement for permutation at practical sample sizes.

**Table S1.** The twenty-one scanned studies of the pre-specified panel, ordered by number of gene-level flips. Columns give the GEO accession, the two-group contrast, the per-arm sample sizes, the number of Hallmark sets with at least fifteen mapped genes, the maximum within-set c^, the number of flips, and the split of those flips into agreeing with permutation (GOOD) versus departing from it (BAD); the split is omitted where there are no flips and is uninformative at n = 3 because permutation cannot declare. The nineteen excluded accessions and their exclusion reasons are listed in the reproducibility deposit (fetchlist_studies.csv).

| **Study** | **Contrast** | **Per arm** | **Sets** | **max c^** | **Flips** | **GOOD/BAD** |
|---|---|---|---|---|---|---|
| GSE302984 | CAF-conditioned vs control medium | 3v3 | 50 | 0.426 | 67 | 60/7 |
| GSE302049 | siCTRL vs siTEDAR | 3v3 | 50 | 0.180 | 49 | 49/0 |
| GSE312436 | control vs minnelide | 3v3 | 30 | 0.892 | 40 | 40/0 |
| GSE131141 | control vs PRDM16 overexpression | 3v3 | 50 | 0.866 | 27 | 24/3 |
| GSE300765 | acidosis 10 weeks vs 48 hours | 3v3 | 50 | 0.082 | 18 | 17/1 |
| GSE303312 | low vs normal glucose | 4v4 | 50 | 0.099 | 13 | 4/9 |
| GSE298981 | IL-22 vs control | 4v4 | 50 | 0.263 | 8 | 2/6 |
| GSE330622 | shCSF2 knockdown vs control | 3v3 | 50 | 0.112 | 4 | 4/0 |
| GSE315735 | carboplatin vs vehicle | 5v4 | 19 | 0.186 | 3 | 2/1 |
| GSE315733 | carboplatin vs vehicle | 5v4 | 24 | 0.191 | 2 | 2/0 |
| GSE315732 | carboplatin vs cisplatin | 3v3 | 24 | 0.437 | 1 | 1/0 |
| GSE313668 | control vs gamma-secretase inhibitor | 3v3 | 50 | 0.362 | 1 | 1/0 |
| GSE312232 | combination vs control | 3v3 | 30 | 0.272 | 1 | 1/0 |
| GSE297053 | PNPase vs SUV3 knockdown | 8v8 | 18 | 0.227 | 1 | 1/0 |
| GSE298668 | LY2090314 vs comparator | 4v4 | 27 | 0.129 | 1 | 1/0 |
| GSE143615 | no vs 4-day tetracycline | 4v4 | 50 | 0.840 | 0 | n/a |
| GSE300351 | CAF19 vs CAF24 | 3v3 | 50 | 0.637 | 0 | n/a |
| GSE230535 | no vs added inhibitor | 4v4 | 50 | 0.409 | 0 | n/a |
| GSE333664 | EHMT1 KO vs EHMT1/2 DKO | 3v3 | 27 | 0.274 | 0 | n/a |
| GSE325279 | DIMATE vs untreated | 4v4 | 34 | 0.177 | 0 | n/a |
| GSE298667 | LY2090314 vs comparator | 4v4 | 21 | 0.096 | 0 | n/a |

**Table S2.** Flip adjudication by per-arm sample size across the twenty-one scanned studies. GOOD counts flips in which the corrected verdict matches the permutation verdict; BAD counts flips in which the naive verdict matches permutation. At n = 3 permutation cannot declare at alpha = 0.05 (ten permutations; minimum p-value 0.1), so its column carries no permutation adjudication; the n = 3 GOOD/BAD entries instead report the flip direction (197 conservative retractions, 11 liberal), whereas at n >= 4 GOOD/BAD report agreement with the permutation gold standard.

| **Per-arm size** | **Studies** | **Flips** | **GOOD** | **BAD (conservative / liberal)** | **Permutation informative** |
|---|---|---|---|---|---|
| n = 3 | 11 | 208 | 197 | 11 | no (structurally powerless) |
| n >= 4 | 10 | 28 | 12 | 16 (12 / 4) | yes |
